# Basement Membrane Structural Integrity Dictates Trans-Tissue Deposition of Laminin in Mammals

**DOI:** 10.1101/2025.04.28.650774

**Authors:** Kohei Omachi, Meei-Hua Lin, Pongpratch Puapatanakul, Karen K. McKee, Hironobu Fujiwara, Peter D. Yurchenco, Jeffrey H. Miner

## Abstract

Basement membranes (BMs) are specialized extracellular matrices (ECMs) essential for tissue structure and function. In non-vertebrates, ECM components can be produced both locally and by distant tissues. In contrast, mammalian ECM has traditionally been considered to originate predominantly from adjacent or tissue-resident cells. The kidney glomerular basement membrane (GBM), composed of laminin-α5β2γ1 and collagen-α3α4α5(IV), is produced by neighboring epithelial cells and functions as a filtration barrier. Alport syndrome, a genetic kidney disease in children, is characterized by GBM structural defects and ectopic laminin-α2 deposition, but the source of this laminin remains unknown. Here, using CRISPR/Cas9 transgenic models, we demonstrated that ectopic laminin-α2 originates not from local kidney cells but from the circulation. Furthermore, laminin-α2 in the mesangium partially derives from circulating sources even under healthy conditions. Our findings uncover a non-cell-autonomous mechanism whereby GBM integrity regulates circulating protein incorporation, revealing a previously unrecognized trans-tissue regulation of BM composition in mammals.

## Introduction

Basement membranes (BMs) are specialized extracellular matrices (ECMs) that play crucial roles in maintaining tissue structure and cell function. Located beneath epithelial cells and surrounding various mesenchymal cells, such as those in muscle and adipose tissue, BMs are composed primarily of laminin, type IV collagen, nidogen, and heparan sulfate proteoglycan. Laminin and type IV collagen form independent polymeric networks bridged by nidogen and other associated molecules ^1,2^.

Laminins are large heterotrimeric glycoproteins composed of α, β, and γ chains. They possess N-terminal polymerization domains that mediate laminin–laminin interactions, long coiled-coil domains that mediate trimerization, and large C-terminal laminin globular (LG) domains that bind cellular receptors to induce signaling. There are at least 16 isoforms in mammals, and each laminin preferentially binds to specific integrin receptors and/or dystroglycan. Laminins play crucial roles in cell differentiation, organogenesis, and homeostasis, in part via tissue-specific expression and localization patterns.

Type IV collagen networks composed of α chain heterotrimers provide mechanical stability to BMs. α chains consist of three domains: an N-terminal 7S domain that mediates trimer-trimer interactions, a triple helical collagenous domain that forms lateral associations ^3–5^ and a non-collagenous (NC1) domain that facilitates both heterotrimerization and trimer-trimer interactions. There are six distinct type IV collagen α chains that assemble into three heterotrimeric isoforms: α1α1α2, α3α4α5, and α5α5α6. The α1α1α2 isoform is expressed ubiquitously, whereas the α3α4α5 and α5α5α6 isoforms display restricted distributions. Dysfunction of type IV collagens and laminins leads to defects in multiple tissues and associated diseases, indicating that this two-polymer network within BMs is essential for development and tissue homeostasis.

In non-vertebrates such as *C. elegans* and *Drosophila*, BM components are produced not only by local cells but also by distant tissues. For example, in post-embryonic *Drosophila*, type IV collagen is synthesized in the fat body and incorporated into BMs throughout the body ^6^. During embryogenesis, hemocytes are major contributors to type IV collagen deposition throughout the body ^7^. In *C. elegans*, muscle-derived type IV collagen is deposited onto distant tissues such as the intestinal BM ^8^. These examples demonstrate a clear trans-tissue mechanism of BM deposition in non-vertebrates. However, whether such a mechanism exists in mammals is unknown.

In mammals, the kidney is one of the organs richest in BMs. In particular, the glomerulus contains a specialized BM —the glomerular basement membrane (GBM)— which is essential for blood filtration. The mature GBM consists of two key molecular networks: laminin α5β2γ1 (LM-521) and type IV collagen α3α4α5 (COL4A3/4/5), which are synthesized by glomerular epithelial podocytes. The GBM interfaces with two specialized cell types, podocytes and fenestrated endothelial cells ^9^, and the health of all three components is critical to the glomerular filtration barrier. Injury to or functional defects in these components disrupt filtration and can lead to severe kidney diseases ^10^.

Alport syndrome is the most common genetic kidney disease in childhood; it is caused by mutations in *COL4A3*, *COL4A4*, or *COL4A5* ^11–13^. In Alport syndrome, the GBM lacks or contains dysfunctional COL4A3/4/5, and its structure and function progressively deteriorate after birth ^14^. Another isoform, COL4A1/1/2, is localized in a thin layer on the endothelial side of the GBM and appears upregulated in Alport syndrome, although it remains confined primarily to the sub-endothelial region ^15^. In addition to these changes in COL4 composition, the laminin repertoire of the GBM is also altered ^16^. While the normal GBM contains only LM-521, in Alport GBM, structurally defective regions also contain ectopic laminin α1 (LAMA1) and laminin α2 (LAMA2) ^17,18^. Notably, the ectopic deposition of LAMA2 is consistently observed across species in Alport GBM and is considered pathogenic. It is proposed that ectopic LAMA2 activates focal adhesion kinase in podocytes, triggering the downstream activation of matrix metalloproteinases and pro-inflammatory cytokines ^19^. Those previous studies concluded that mesangial cells, which normally express LAMA2, are its source for the Alport GBM, but it is unclear whether they are the sole contributors. Given the high renal blood flow, one hypothesis is that LAMA2 synthesized in distant tissues enters the circulation and is subsequently incorporated into the structurally compromised regions of GBM.

To test this hypothesis, we investigated the association between ectopic LAMA2 deposition and GBM structural defects and found primary GBM abnormalities contribute to LAMA2 deposition. To further examine the origin of LAMA2, we generated two in vivo CRISPR/Cas9 mouse models; one used live Cas9 ^20^ to mutagenize *Lama2*, and the second used dead (d)Cas9-KRAB ^21^ to repress *Lama2* expression. Comparing these models revealed that Cas9-mediated knockout was more efficient in eliminating LAMA2 deposition than the dCas9-KRAB repression approach. Cell-specific *Lama2* knockout studies revealed that ectopic LAMA2 originates from tissues outside the kidney. Interestingly, human and mouse plasma contain LAMA2 protein, indicating that LAMA2-containing laminins circulate in the blood. Moreover, intravenous (i.v.) injections of recombinant LM-211 protein into *Lama2^-/-^* mice showed that circulating LAMA2 can be deposited into the abnormal Alport GBM, as well as into the glomerular mesangium of both healthy and Alport mice. Together, these findings uncover a non-cell-autonomous mechanism in which the GBM’s structural integrity dictates its incorporation of circulating LAMA2. Furthermore, we show that a portion of circulating LAMA2 is incorporated into the mesangial matrix under physiologic conditions. Our data reveal that, like in non-vertebrates, mammalian ECM exhibits a trans-tissue deposition mechanism in both health and disease.

## Results

### Laminin α2 is specifically deposited in structurally defective GBMs

Although LM-521 is the predominant isoform in the mature GBM, ectopic deposition of laminin containing the laminin α2 chain (LAMA2) is observed in Alport GBM. Laminin trimers containing LAMA2 are normally located in the glomerular mesangial matrix, a non-BM matrix ^18^. Abnormal LAMA2 deposition in the GBM has been proposed to be pathogenic, either directly or indirectly perturbing podocyte homeostasis ^19,22^. To determine whether ectopic GBM LAMA2 deposition is a common feature across various glomerular diseases, we compared four glomerular disease models with defects in the filtration barrier. These included two primary GBM diseases, Alport syndrome (*Col4a5* knockout) ^23^ and Pierson syndrome (*Lamb2* knockout) ^24^, as well as two non-GBM-associated diseases: the *Cd2ap* knockout ^25^ and Adriamycin nephropathy (ADRN) ^26,27^. Pierson syndrome is caused by mutations in *LAMB2*, which encodes laminin β2, a subunit of GBM LM-521. Loss of LAMB2 results in GBM defects and nephrotic syndrome. CD2AP functions as an adapter protein in podocyte foot process, where it binds to the cytoplasmic domain of nephrin, a key component of the slit diaphragm. Knockout of *Cd2ap* induces effacement of podocyte foot processes, leading to glomerular filtration defects. Adriamycin is used experimentally to induce glomerular disease characterized by disruption of the podocyte cytoskeleton. Given these distinct disease mechanisms, both Alport syndrome and Pierson syndrome result in a structurally defective GBM, whereas the *Cd2ap* knockout and ADRN models cause podocyte injury and foot process abnormalities without affecting the GBM’s composition or ultrastructure.

Immunofluorescence analysis revealed that the mesangial matrix contained LAMA2 in both healthy and diseased conditions (Figure 1A-Η). Ectopic LAMA2 deposition in the GBM was observed in the two primary GBM disease models (Figure 1Β, D) but was absent in the primary podocyte disease models (Figure 1F, H). These results indicate that ectopic LAMA2 deposition is a feature associated with a GBM structural defect rather than with any glomerular defect causing proteinuria. Notably, while the LAMA2-containing mesangial matrix was expanded in both *Lamb2* and *Cd2ap* knockout mice, LAMA2 was absent from the *Cd2ap* knockout GBM. This suggests that mesangial matrix expansion is unrelated to the ectopic LAMA2 deposition in the GBM.

**Figure. 1.**
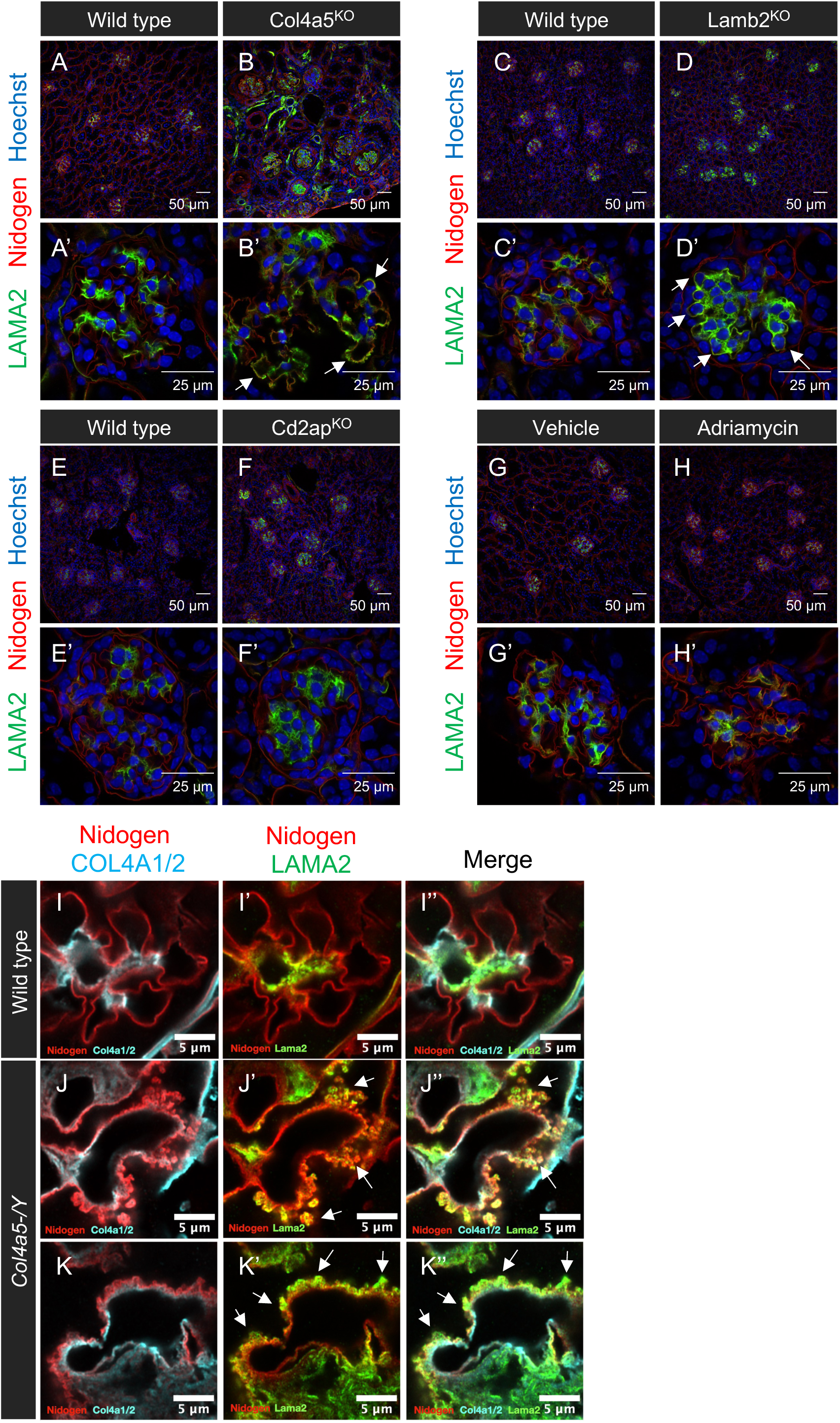
Laminin α2 protein is deposited in structurally defective GBMs. Immunofluorescence was used to localize LAMA2 (green) and Nidogen (red) in kidney sections. (A, D) Laminin α2 was deposited in the defective GBMs (arrows) of *Col4a5* mutant and *Lamb2* mutant mouse models of Alport and Pierson syndromes, respectively. (E, F) LAMA2 localization was restricted to the mesangial matrix in the *Cd2ap* mutant, despite mesangial matrix expansion. (G, H) LAMA2 was localized to the mesangial matrix in the Adriamycin-treated mouse, a pattern similar to control. (I-K) Airyscan imaging showed the ectopic LAMA2 (green) deposited on top of the COL4A1/2 (red) layer, overlapping with the split GBM labeled by nidogen, which marks the podocyte side of the GBM.

The peripheral capillary loop GBM that mediates filtration interfaces with two cell types: podocytes and endothelial cells. To further investigate the specific location of ectopic LAMA2 within the GBM, we examined its localization in Alport mice using near super-resolution microscopy. Consistent with Figure 1B, healthy GBM did not contain LAMA2, whereas Alport GBM did. In healthy glomeruli, COL4A1/2 is in a thin layer at the endothelial aspect of the GBM. In Alport GBM, COL4A1/2 appears increased in response to the loss of COL4A3/4/5, yet it remains predominantly localized to the endothelial side ^15^. By comparing COL4A1/2, nidogen, and LAMA2 in Alport GBM, we observed that LAMA2 was positioned distal to COL4A1/2 and overlapped with split GBM labeled by nidogen, indicating that the ectopic LAMA2 was deposited on the podocyte aspect of the Alport GBM (Figure 1I-K). A previous report proposed that mesangial cell processes extend into the peripheral capillary loop GBM and deposit LAMA2 into the Alport GBM ^22^. While it is reasonable to suggest that mesangial cells are the source of ectopic LAMA2, given that they are the only glomerular cells known to express LAMA2, there is evidence against this ^28^. Our findings show that LAMA2 is located on the podocyte side of the peripheral GBM, far from the mesangium. Moreover, mesangial expansion was not associated with ectopic LAMA2 deposition. Based on these observations, we hypothesized that there is a different source of LAMA2 in Alport GBM.

### Circulating laminin α2 as a potential source for GBM deposition

Given the high blood flow to the kidneys, circulating proteins have ready access to renal tissues. In particular, the glomerulus experiences robust vascularization to facilitate blood filtration, making the circulation a potential source of ectopic LAMA2. To investigate whether LAMA2 is normally in the blood, we first determined the repertoire of ECM proteins in human plasma using PeptideAtlas, a publicly accessible proteome database ^29–31^. We searched for major laminins and collagens, as well as fibronectin, a well-known circulating ECM protein ^32^ that is also present ectopically in Alport GBM ^33^. Numerous peptides derived from ECM proteins such as LAMA2, LAMB1, LAMC1, COL1A1, COL5A1, COL6A, COL18A1, and fibronectin were identified in human plasma. Interestingly, LAMA2, LAMB1, and LAMC1, which form the Laminin-211 (LM-211) heterotrimer, were abundant in plasma, whereas other laminin proteins were detected at low levels or not at all (Figure 2A, Figure S1, Table S1). Protein coverage for LAMA2, LAMB1, and LAMC1 were 70.3%, 77.8%, and 76.1% respectively (Figure S2).

**Figure 2.**
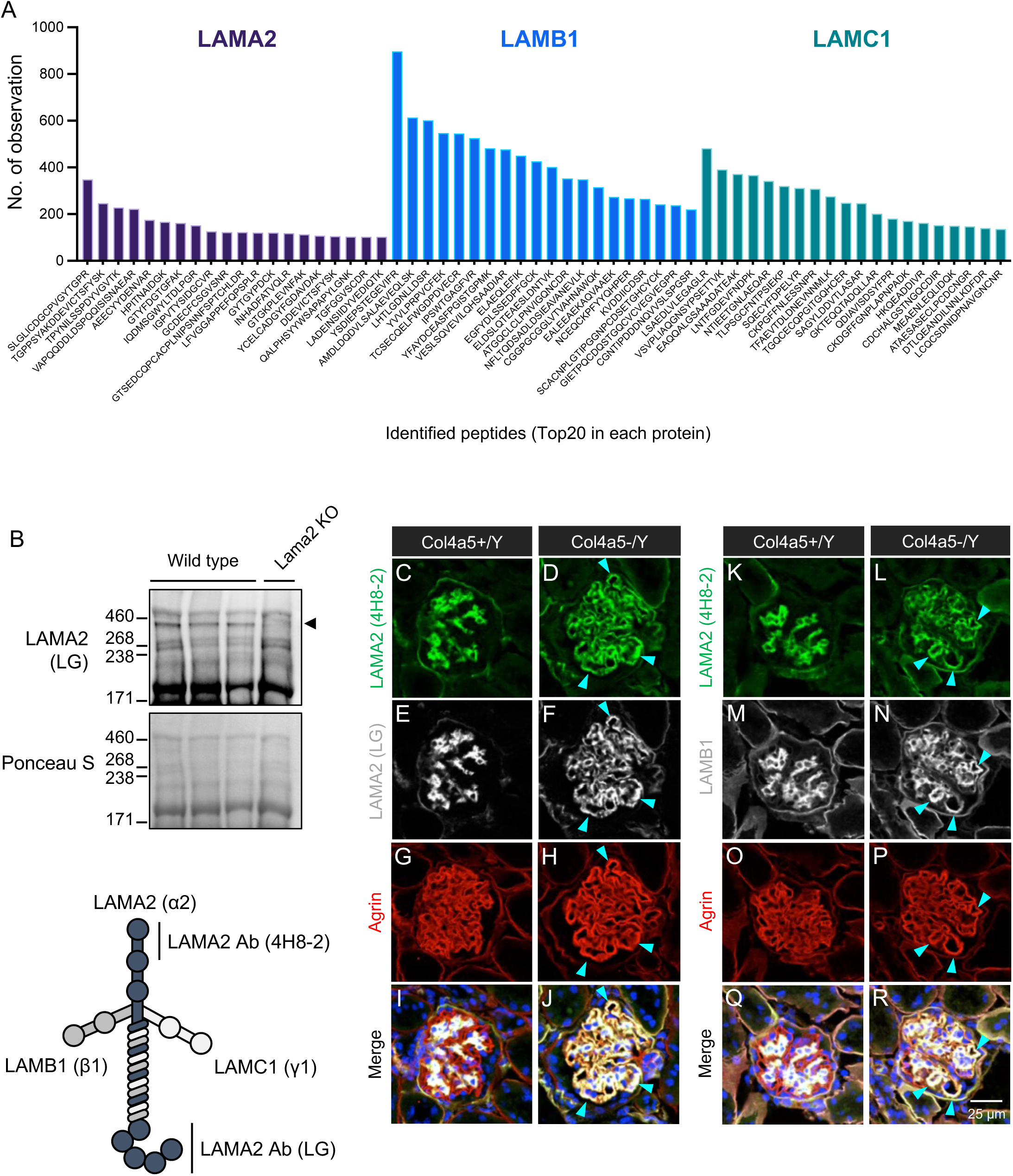
Laminin α2/LM-211 circulates in the blood in both humans and mice. (A) The number of observations for each unique peptide derived from LAMA2, LAMB1, and LAMC1 in human plasma. Over one hundred peptides related to LM-211 were identified in human plasma, indicating that LAMA2 circulates in the blood. (B) Immunoblotting using anti-LAMA2 antibody recognizing the C- terminal LG domains confirmed its presence in mouse plasma. The LAMA2 signal observed in wild-type plasma was absent in *Lama2* knockout (dy3k/dy3k) mice (arrowhead). Ponceau S stain was used for visualizing total protein on the membrane. (C-J) Immunostaining for LAMA2 N-terminus (4H8-2) and C- terminus (LG), along with agrin (GBM), demonstrated that ectopic LAMA2 was likely intact, with both ends present (arrowheads). (K-R) Immunostaining for LAMA2 N-terminus (4H8-2) with LAMB1 (a component of LM-211) and agrin (GBM) revealed that ectopic LAMA2 colocalized with LAMB1 (arrowheads).

We next focused on the mouse by carrying out western blotting of plasma from wild-type mice, which detected LAMA2 signals that were absent in *Lama2* knockout (dy3k/dy3k) mice (Figure 2B). The molecular weight of LAMA2 corresponded to its full-length form. These data strongly support the presence of LAMA2 in the circulation. Moreover, immunofluorescence analysis revealed that the ectopic LAMA2 in the GBM contained both N- and C-terminal regions (Figure 2C–J) and colocalized with LAMB1, a component of LM-211 (Figure 2K–R), consistent with the concept that LAMA2 was not fragmented and was likely complexed with LAMB1. These results support our hypothesis that the circulation is a potential source of ectopic LAMA2.

### Development of CRISPR/Cas9 knockout and repression mice to block LAMA2 expression

To investigate the origin of ectopic LAMA2 in vivo, we employed CRISPR/Cas9-mediated knockout and transcriptional interference in mice. We first screened active single guide RNAs (sgRNAs) for *Lama2* knockout and transcriptional repression in cultured cells. For *Lama2* knockout, sgRNAs targeting protein-coding exons were selected. For transcriptional repression, sgRNAs targeting regions near the transcription start site (TSS) were chosen (Figure 3A). T7 endonuclease 1 (T7E1)-mediated mismatch cleavage assays revealed that eleven sgRNAs were effective in introducing insertions and deletions (indels) in Cas9-expressing Neuro-2a cells (Figure 3B and Figure S3A). sgRNAs for *Lama2* knockdown were then tested in dCas9-KRAB-expressing C2C12 cells. Quantitative RT-PCR demonstrated that four sgRNAs significantly repressed *Lama2* transcription (Figure 3C and Figure S3B). We selected three (#1 (=#3’), #2 and #3) to construct a single transgene (Tg) used to generate Lama2-gRNA transgenic mice (Tg: Lama2-gRNA, Figure S4). Because one of these sgRNAs (#1) targets the coding region in exon 1, Tg: Lama2-gRNA can be used in both Cas9 and dCas9-KRAB mice, allowing us to directly compare their efficacies.

**Figure 3.**
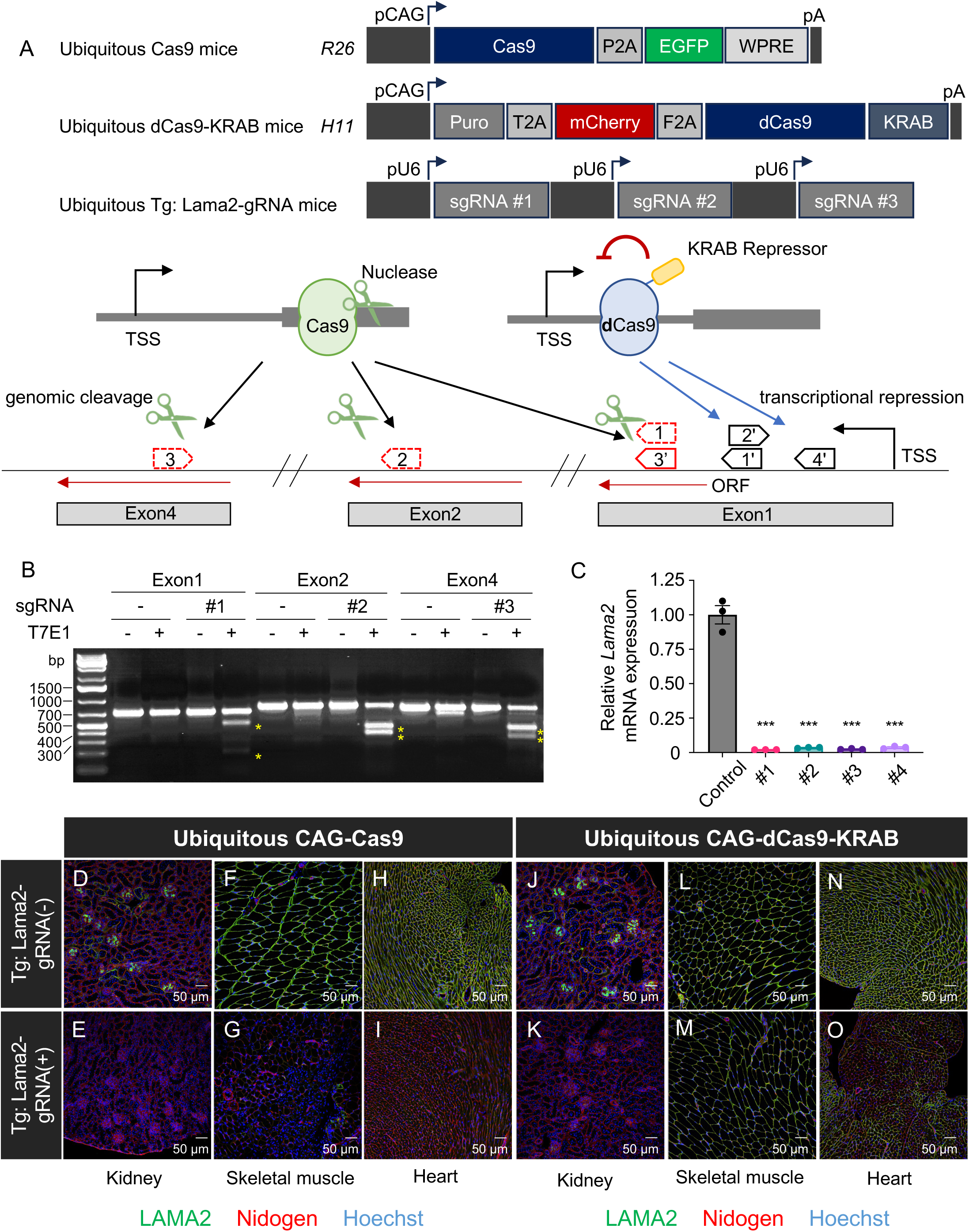
In vivo CRISPR-KO and CRISPRi blocked Laminin α2 protein expression. (A) The schematic diagram illustrates the Cas9, dCas9-KRAB, and Tg: Lama2-gRNA constructs, as well as the gRNA targeting sites within the mouse *Lama2* gene. In the CRISPR-KO approach, Cas9 induces double-strand breaks, leading to indel formation within the protein-coding sequence. In contrast, CRISPRi utilizes dCas9-KRAB to suppress transcription near the transcription start site (TSS). The red-highlighted sgRNAs were selected for inclusion in the transgene. (B) T7E1 assay showed mismatch cleavage activity in amplicons from Cas9 expressing Neuro-2a cells (yellow asterisks). All three sgRNAs introduced indels. (C) Expression analysis of endogenous *Lama2* in C2C12 cells. All four sgRNAs significantly decreased *Lama2* expression. Error bars indicate the mean ± SE (n=3). Statistical analysis was performed using one-way ANOVA with Dunnett’s multiple comparisons test. ***, P <0.001 vs. control. (D-F) Direct comparison of CRISPR-KO and CRISPRi in various tissues expressing LAMA2 protein by immunofluorescence. Ubiquitous Cas9 (Rosa26-CAG-Cas9); Tg: Lama2 gRNA mice showed greatly decreased LAMA2 in kidney (D, F), skeletal muscle (F, G), and heart (H, I). Ubiquitous dCas9-KRAB (H11-CAG-dCas9-KRAB); Tg: Lama2 gRNA mice showed greatly decreased LAMA2 in kidney (J, K) but only partial reduction in skeletal muscle (L, M) and heart (N, O).

To test the functionality of the gRNAs in vivo, we crossed the Tg: Lama2-gRNA mice with ubiquitous CAG-Cas9 and CAG-dCas9-KRAB mice. Immunofluorescence analysis revealed that both the CAG-Cas9; Tg: Lama2-gRNA and CAG-dCas9-KRAB; Tg: Lama2-gRNA models exhibited significantly reduced LAMA2 deposition throughout the kidney (Figure 3D, E, J, K). However, the reduction of LAMA2 in skeletal muscle and heart was more pronounced in the CAG-Cas9 model compared to the CAG-dCas9-KRAB model (Figure 3F-I and L-O). Given that *Lama2* expression is much higher in skeletal muscle and heart than in kidney, it is likely that although dCas9-KRAB strongly reduced Lama2 mRNA levels in these tissues, residual mRNA was sufficient to produce LAMA2 protein, leading to its accumulation. In contrast, Cas9-mediated indels should prevent the production of functional LAMA2 protein. Indeed, ubiquitous *Lama2* knockout by Cas9 caused severe muscular dystrophy in these mice (Figure S5), which is similar to the phenotype observed in *Lama2* knockout mice ^34^. However, ubiquitous *Lama2* knockdown by dCas9-KRAB did not result in severe muscular dystrophy, but did cause hind limb abnormalities, including paralysis (data not shown). Based on these findings, a Cas9-mediated knockout approach appears more effective for targeting proteins with long half-lives, such as ECM proteins.

### Ectopically deposited LAMA2 is not derived from glomerular cells

We next utilized Cre-mediated activation of Cas9 expression in a tissue specific fashion to investigate the source of ectopic LAMA2 in the GBM. *Pax3* is a paired homeobox gene expressed in early kidney progenitors that give rise to the entire nephron and to the stromal compartment. Based on extensive lineage tracing using the Pax3-Cre transgene, large subsets of cells in the mature kidney, including podocytes, mesangial cells, endothelial cells, and interstitial cells originate from cells that expressed Cre at some point ^35–37^. Therefore, the Pax3-Cre transgene allows the activation of Cre-dependent genes in almost all glomerular constituent cells and in stromal cells. Taking advantage of the specific yet broad glomerular cell-targeting properties of Pax3-Cre, we knocked out *Lama2* in glomerular and stromal cells by generating Pax3-Cre; CAG-LSL-Cas9; Tg: Lama2-sgRNA mice. Immunofluorescence analysis revealed that *Lama2* knockout in Pax3-Cre lineage cells significantly reduced LAMA2 protein in the kidney interstitium. Surprisingly, although LAMA2 in the mesangial matrix was slightly decreased, it remained detectable (Figure 4A, B). Given the activity of Pax3Cre in mesangial and other glomerular cell precursors observed with the Ai14 Cre reporter (Figure S6A), the *Lama2* gene should be disrupted in mesangial cells. Indeed, Lama2 mRNA was disrupted in kidney cortical cells (Figure S7). However, LAMA2 protein was still detected, suggesting that part of the LAMA2 in the mesangial matrix may be supplied by non-Pax3-Cre lineage cells. We revisit this result in more detail below.

**Figure 4.**
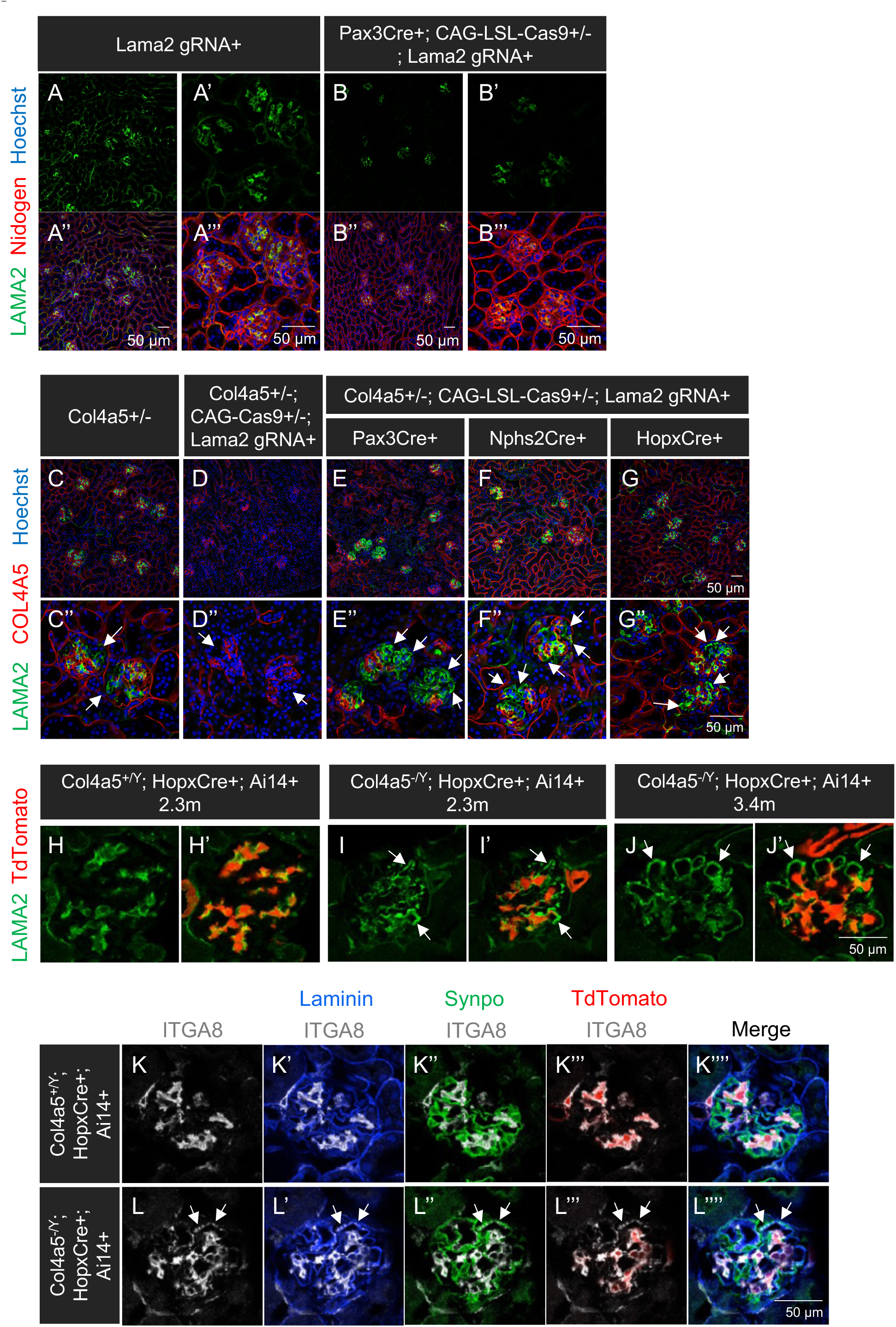
Ectopically deposited Laminin α2 in Alport mouse GBM is not derived from glomerular cells. (A, B) Immunostaining of LAMA2 and nidogen. Pax3Cre; CAG-LSL-Cas9; Tg: Lama2 gRNA mice showed strongly reduced LAMA2 in the interstitium, but only modest reduction in the mesangial matrix. This suggests that some of the mesangium’s LAMA2 is derived from external tissues. (C-G) Ubiquitous Cas9; Tg: Lama2 gRNA mice showed no LAMA2 in COL4A5-negative GBM segments in female *Col4a5^+/-^* mice (arrows vs arrowheads). However, Pax3-Cre- (include podocytes, mesangial cells and interstitial fibroblasts) and Podocin-Cre- (podocytes) dependent Cas9; Tg: Lama2 gRNA mice showed little reduction in ectopic LAMA2 GBM deposition (arrows). This suggests that the ectopic LAMA2 is derived from external tissues. (H-J) HopxCre-tdTomato reporter mice showed that there were no mesangial cell processes in the LAMA2-positive GBM segments in Alport mice (arrows). (K, L) Immunostaining for ITGA8 along with anti-Synpo (podocytes), anti-laminin (GBM), and tdTomato expression (mesangial cells) demonstrated that ITGA8 is present on Alport but not WT podocytes. As in H-J, no mesangial cell processes (red) were observed in WT or Alport peripheral capillary loop GBMs.

Next, we used Pax3-Cre activation of Cas9 to knockout *Lama2* in Alport mice to determine whether the ectopic LAMA2 originates from glomerular cells. For this purpose, we used female *Col4a5* heterozygous mice. Because *Col4a5* is on the X chromosome, and one of the two *Col4a5* alleles (wild-type or null) is randomly inactivated in podocyte precursors, *Col4a5^+/-^* females have both normal (COL4A3/4/5-positive) and mutant (Alport COL4A3/4/5-negative) GBM segments in the same glomeruli (Figure 4C), allowing us to directly compare adjacent healthy and Alport GBM. Utilizing this model, we investigated whether Pax3-Cre-driven *Lama2* knockout (which occurs independently of X chromosome inactivation) blocks ectopic LAMA2 deposition in Alport GBM. Although ubiquitous Cas9-mediated *Lama2* knockout diminished LAMA2 throughout the kidney (Figure 4D), Pax3-Cre/Cas9-mediated *Lama2* knockout reduced LAMA2 in the interstitium but not in the mesangial matrix nor in the Alport GBM segments (Figure 4E). Furthermore, neither podocyte-specific Cre (Nphs2-Cre) nor mesangial-specific Cre (Hopx-Cre, Figure S4B) was effective in blocking the deposition of ectopic LAMA2 in the Alport GBM (Figure 4F, G). These targeted genetic approaches suggest that the ectopic LAMA2 is not derived from Pax3-lineage resident glomerular cells, including mesangial cells and podocytes.

Previous studies have proposed that mesangial cell processes extend far into the Alport peripheral capillary loop GBM and deposit LAMA2 into the injured GBM ^19,22,38^. To investigate this possibility, we examined whether mesangial cell processes can be detected in the Alport GBM using Hopx-CreER; Ai14 and Hopx-Cre; Ai14 to label mesangial cells by forcing them to express tdTomato. We confirmed that mesangial cell bodies were efficiently labeled with tdTomato in Hopx-CreER; Ai14 mice injected with tamoxifen and in HopxCre; Ai14 mice (Figure S6B, C). Although the mesangium appeared expanded in some glomeruli in hemizygous male *Col4a5^-/Y^* Alport; Hopx-Cre; Ai14 mice and *Col4a5^-/Y^* Alport; Hopx-CreER; Ai14 mice, there were no tdTomato-positive processes in the peripheral capillary loop GBM (Figure 4H-J and Figure S8).

Integrin α8 is a well-known mesangial cell marker ^39^ and a receptor for nephronectin in the GBM ^40^. Previous studies concluded that cell processes in the peripheral Alport GBM were mesangial cell-derived based in part on positive integrin α8 immunostaining near the peripheral capillary loop GBM. Here, Hopx-Cre; Ai14-labeled mesangial cells were found adjacent to integrin α8-positive capillary loops in Alport mice, but no tdTomato was observed in the peripheral loops (Figure 4K, L). Furthermore, high-resolution confocal microscopy revealed that integrin α8 was localized basally on podocytes above the basement membrane and overlapping with synaptopodin (Figure 4L’’), a podocyte foot process cytoskeletal protein. This indicates that the integrin α8-positive cells adjacent to the Alport GBM are podocytes, not mesangial (or endothelial) cells. Overall, these results show that the ectopic LAMA2 in the Alport GBM is not synthesized by glomerular cells. Moreover, the presence of LAMA2 in the mesangium of Pax3-Cre/Cas9-mediated knockout mice (Figure 4A) suggests that at least some of the mesangial LAMA2 is not synthesized by mesangial cells.

### Circulating laminin α2 is non-cell autonomously deposited in structurally defective GBMs

Since mesangial cells and interstitial fibroblasts express *Lama2* but are unlikely to be the major source of ectopic LAMA2, we hypothesized that laminin trimers containing LAMA2 (e.g., LM-211) circulate in the blood and are gradually deposited into any “gaps” in the defective GBM. To investigate the hypothesis that circulating LAMA2 can accumulate in the defective Alport GBM, we intravenously injected recombinant human (h)LM-211 protein into both normal and Alport mice with ubiquitous Cas9-mediated *Lama2* knockout. As shown in Figure 1*A*, structural defects in the GBM are associated with ectopic LAMA2 deposition. The intravenously injected hLM-211 reached the kidney and deposited in the mesangial matrix in both healthy and Alport mice (Figure 5A-D), but in addition, it was deposited into the GBM of Alport mice (Figure 5D, F). Furthermore, in *Col4a5^+/-^* female mice, the injected hLM-211 was specifically deposited in the COL4A5-negative GBM segments (Figure 5G, H). Taken together, our data reveal that a fraction of LAMA2 in the mesangial matrix and all the LAMA2 in the Alport GBM originate from the circulation rather than from glomerular cells. This mode of BM protein deposition has not been previously reported, highlighting the existence of the trans-tissue deposition mechanism for BM protein in mammals.

**Figure 5.**
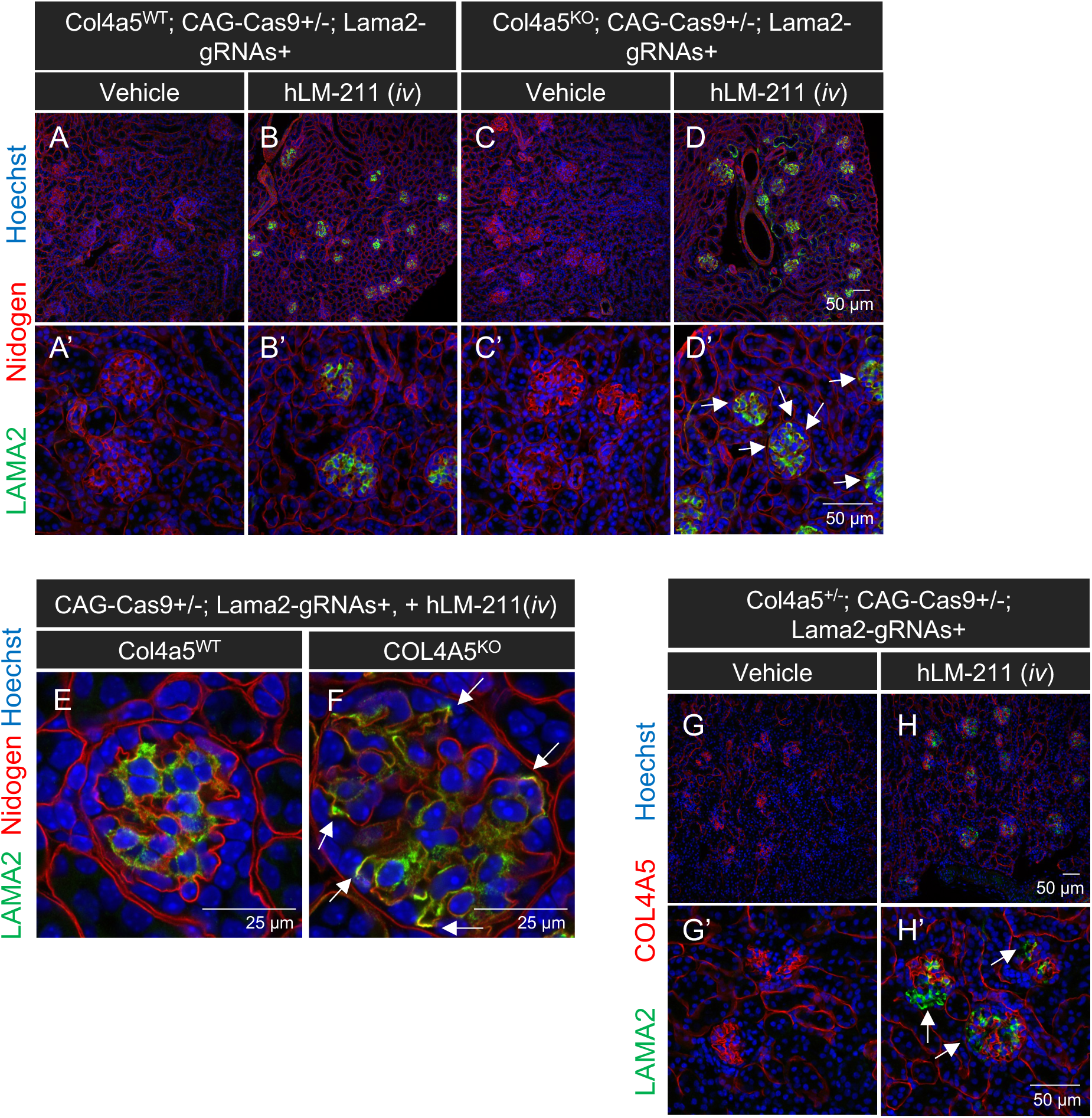
Circulating Laminin α2/LM-211 was deposited in the mesangial matrix in *Lama2* knockout *Col4a5* WT glomeruli and in the injured GBM in *Lama2* knockout *Col4a5* mutant mice. (A-D) Localization of i.v. administered recombinant LM-211 in the kidneys of LAMA2-deficient *Col4a5* WT and mutant mice. Circulating LM-211 protein was deposited in the mesangial matrix and in Alport GBM (arrows). (E, F) Higher magnification images show that injected recombinant LM-211 was deposited in the structurally defective GBM in Alport glomeruli (arrows), mimicking the ectopic LAMA2 deposition observed in Alport mice. (G, H) Localization of i.v. administered recombinant hLM-211 in the kidney of LAMA2-deficient *Col4a5*+/- mice. LM-211 was specifically deposited in GBM segments lacking COL4A5 (arrows).

## Discussion

In non-vertebrates such as *C. elegans* and *Drosophila*, it is well established that ECM proteins are synthesized in both local and distant tissues and deposited into BMs. The latter exemplifies a trans-tissue mechanism of BM incorporation. In mammals, however, such a mechanism has not been reported. In this study, we identified a trans-tissue deposition of circulating laminin-α2 (LAMA2) into the structurally defective glomerular BM (GBM) of the mouse kidney. Interestingly, LAMA2 is present only in structurally compromised GBMs, indicating that this trans-tissue mechanism is regulated by BM structural integrity. Furthermore, beyond the GBM, circulating LAMA2 is also partially incorporated into the mesangial matrix even under healthy conditions. These findings suggest that trans-tissue BM deposition exists in mammals.

Although ectopic deposition of laminins in Alport GBM was reported over 20 years ago ^18^ and was proposed to be pathogenic ^19^, the origin of these laminins has remained unclear. In this study, we used a transgenic CRISPR/Cas9 approach to eliminate LAMA2 from the Alport GBM in order to investigate its source. Our findings reveal a previously unrecognized mechanism that ectopic LAMA2 in Alport GBM does not originate from local glomerular cells. Our data suggest that LAMA2 circulates in the blood as part of the LM-211 trimer and is selectively deposited in the structurally defective Alport GBM but not in normal GBM. Circulating LAMA2 was also deposited in the mesangial matrix, a matrix known to accumulate plasma proteins ^41^. We propose a non-cell-autonomous mechanism in which the integrity of the glomerular filter determines whether circulating LAMA2 is deposited in the GBM. Specifically, we suggest that an intact GBM is a highly selective barrier that excludes the entry of the ∼700 kDa LM-211 trimer and also lacks the necessary space for its deposition, whereas a defective GBM loses structural integrity and organization, creating larger pores as well as the space needed for LM-211 deposition. This mechanism could explain why ectopic LAMA2 is observed exclusively in structurally defective GBMs. Notably, the ectopic LAMA2 likely originates from outside the kidney. The source of LAMA2 in Alport GBM has been a subject of controversy for many years, making this finding particularly important.

Mesangial cells, which are smooth muscle-like contractile cells ^42^, normally express *Lama2*, and LAMA2 is typically restricted to the mesangial matrix under healthy conditions ^43^. Consequently, it was proposed that LAMA2 in Alport GBM is deposited by mesangial cell processes that invade the peripheral capillary loop; this was based on the use of integrin α8 antibody as a specific marker for mesangial cells. However, electron microscopic analysis of serial sections revealed that the processes in Alport GBM are derived from podocytes ^28^. Our results explain this discrepancy by demonstrating that Alport podocytes ectopically express integrin α8. As integrin α8β1 is a receptor for nephronectin in the GBM, their interaction could provide pathogenic signals to podocytes and promote disease progression. Additionally, we labeled mesangial cells in Alport mice with tdTomato using Hopx-CreER; Ai14 and HopxCre; Ai14 but did not see tdTomato in the peripheral capillary loops, indicating there were no mesangial cell processes there. Mesangial cells have been proposed to migrate into Alport GBM, but our findings demonstrated they do not.

To investigate the origin of LAMA2, we employed LAMA2 knockdown in mice using two different in vivo CRISPR/Cas9 systems: Cas9-mediated knockout and dCas9-KRAB-mediated transcriptional repression. Both approaches successfully blocked LAMA2 expression in the kidney, but the Cas9 approach was more effective in skeletal muscle and heart, where LAMA2 is abundantly expressed, compared to dCas9-KRAB. However, of the various cell type-specific knockouts of *Lama2* that we generated, even the broadest one using Pax3-Cre failed to eliminate LAMA2 from the mesangium or from Alport GBM. These results suggested that the ectopic LAMA2 did not originate from kidney cells.

Given these new findings and the substantial volume of blood that passes through the glomeruli, our hypothesis—that ectopic LAMA2 originates from external tissues via the bloodstream—has strong support. Further supporting this hypothesis, specific peptides from LAMA2, as well as from LAMB1 and LAMC1 with which it trimerizes to make LM-211 ^44^, are present in human plasma ^29,31^. Moreover, western blot analysis revealed the presence of the LAMA2 protein in mouse plasma. These indicate that LM-211 circulates in the blood under healthy conditions. The likely source is the heart and/or skeletal muscle, as these tissues are rich in LM-211 and are subjected to physical stresses ^45^, likely leading to LM-211 being shed into the circulation. Finally, we demonstrated that intravenously injected hLM-211 reached the glomerulus and was deposited in the Alport GBM.

Besides LAMA2, the Alport GBM contains other ectopic ECM molecules, including LAMA1 ^17^, fibronectin ^33^, type V collagen, and type VI collagen ^46^, which are not typically found in the normal GBM. LAMA1 has been observed only in mice and has been shown through transmission electron microscopy to be synthesized by podocytes, and LAMA1 is specifically deposited in structurally defective GBM ^17^. This suggests that LAMA2 and LAMA1 in the Alport GBM have different mechanisms of deposition. The origins of types V and VI collagens in the Alport GBM have not been investigated. However, both collagens have been identified in human plasma, similar to LAMA2. Interestingly, type VI collagen is abundant in muscle ^47^, suggesting that its ectopic deposition in the Alport GBM may occur through a mechanism similar to that of LAMA2. Although the deposition of type XIII collagen has not been examined in Alport syndrome, its accumulation was elevated in a mouse model of anti-GBM disease, leading to GBM structural defects due to an acute immune reaction ^48^. These observations support the idea that circulating BM proteins are preferentially deposited into structurally defective GBMs. Beyond the disease context, the mesangial matrix in healthy kidneys contains LAMA2, LAMB1, LAMC1, COL1A1, COL5A1, COL6A1, and COL18A1, all of which were found in human plasma by proteomics (Figure S1). This suggests the existence of trans-tissue ECM regulation even under healthy conditions.

There are limitations to our study. While our data demonstrated that circulating LAMA2 is deposited into structurally defective GBM, the specific tissues contributing to circulating LAMA2 remain unclear. However, given that LAMA2 is highly expressed in cardiac and skeletal muscles, these tissues are likely sources. In addition, we were unable to determine the impact of ectopic LAMA2 on the progression of Alport syndrome. Because the ectopic LAMA2 is deposited through the bloodstream, ubiquitous *Lama2* knockout is required to remove LAMA2 from the Alport GBM. However, this results in severe lethal muscular dystrophy ^34^ (Figure S5), which precludes an analysis of kidney disease progression in *Lama2* knockout Alport mice. Despite this limitation, our findings that BM selectively incorporates circulating proteins suggest a novel therapeutic strategy for restoring BM structure in disease contexts, as discussed below.

From a therapeutic perspective, our findings have the potential to contribute to the development of new treatment strategies for Alport syndrome. BM defects may be repairable through the circulation of proteins in the bloodstream. Indeed, intravenous injection of full-length LM-521 protein temporarily rescues GBM defects in a mouse model of Pierson syndrome caused by a *Lamb2* mutation ^49^. Looking beyond the kidney, in LAMA2-deficient muscular dystrophy, LM-111 delivery through the bloodstream rescues the dystrophic phenotype ^50^. Moreover, genetic introduction of the rationally designed proteins mini-agrin and αLNNd into *Lama2* mutants cooperatively restores BM function by promoting laminin polymerization and interacting with cell surface receptors ^51–53^. These studies strongly support the idea that BM defects may be repairable. To rescue Alport syndrome, protein therapy using type IV collagen itself or rationally designed proteins delivered via the circulation could restore GBM integrity. For the latter approach, a deeper understanding of how BM defects gradually develop during disease progression, as well as the structural and functional differences among type IV collagen isoforms, is essential.

Finally, in the basic science kidney research field, gene targeting in mesangial cells has historically been challenging due to the limited availability of cell-specific Cre lines. While Foxd1-Cre has been used for mesangial cell targeting, its expression in stromal progenitors leads to broad targeting ^54,55^. Recent single-nucleus RNA sequencing data indicates that *Hopx* is specifically expressed in mesangial cells and juxtaglomerular cells ^56–58^, suggesting that Hopx-Cre is a more effective tool for targeting mesangial cells. To explore this, we generated Hopx-Cre/Hopx-CreER; Ai14 mice and demonstrated that both Cre lines effectively labeled mesangial cells and juxtaglomerular cells, providing more specific targeting in the kidney compared to Foxd1-Cre. Additionally, we reexamined Pax3-Cre transgene expression with Ai14 as a reporter and confirmed that Pax3-Cre targets almost all kidney cells except for collecting duct cells of the ureteric bud lineage. With its broad targeting properties, Pax3-Cre is a powerful tool for multi-cell targeting in the kidney. Our study offers a new perspective on mesangial cell function and its role in kidney biology.

In summary, our study revealed a non-cell-autonomous mechanism in which LAMA2 is deposited in structurally defective GBMs, and likely also in the normal mesangium, from the circulation. Developmental roles for a similar process have been demonstrated in *C. elegans* ^8^ and *Drosophila* ^6^. However, ECM in mammals has traditionally been thought to be produced primarily by adjacent cells or by cells within the same tissue. Our study introduces the concept of trans-tissue ECM deposition in mammals, a model that has not been thoroughly investigated but may play an important role in both developmental and disease contexts.

## Materials and Methods

### Plasmid DNA

lentiCRISPR v2 was a gift from Feng Zhang (Addgene plasmid # 52961; http://n2t.net/addgene:52961; RRID:Addgene_52961) ^59^. lentiGuide-Puro was a gift from Feng Zhang (Addgene plasmid # 52963; http://n2t.net/addgene:52963; RRID:Addgene_52963). Lenti-EF1alpha-dCas9-KRAB_Hygro (Addgene plasmid # 192663; http://n2t.net/addgene:192663; RRID:Addgene_192663) and Lenti-EF1alpha-dCas9-KRAB-MeCP2_Hygro (Addgene plasmid # 192664; http://n2t.net/addgene:192664; RRID:Addgene_192664) were generated by Gibson assembly. Briefly, dCas9-KRAB or dCas9-KRAB-MeCP2 were amplified with 20 bp overlapping ends from pB-CAGGS-dCas9-KRAB-MeCP2 (a gift from Alejandro Chavez & George Church, Addgene plasmid #110824; http://n2t.net/addgene:110824; RRID). The lenti MS2-P65-HSF1_Hygro plasmid (a gift from Feng Zhang, Addgene plasmid #61426; http://n2t.net/addgene:61426; RRID) was used as a backbone, amplified for the insertion of dCas9-KRAB or dCas9-KRAB-MeCP2 between the EF1a promoter and T2A-Hygro sites. These amplicons were purified using the QIAquick PCR Purification Kit (QIAGEN, #28106). Purified fragments were then mixed with NEBuilder® HiFi DNA Assembly Master Mix (NEB, #E2621S) and incubated at 50°C for 1 hour. The Gibson assembly product was transformed into NEB® 5-alpha Competent E. coli (NEB, C2987H). Correct plasmids were screened by restriction enzyme digestion and confirmed by Sanger sequencing. The sgRNAs were cloned using Golden Gate assembly method, as previously described. sgRNA sequences are listed in Table *S2* and *S3*. All plasmids used in this study were purified using the PureLink™ HiPure Plasmid Midiprep Kit (Invitrogen, #K210005) by following to the manufacturer’s instructions.

### Recombinant Laminin-211 protein

Recombinant LM-211 was purified as previously described ^60^. Briefly, HEK293 cell lines stably expressing recombinant LM-211 were grown in the presences of furin inhibitor 1 (5 µM in medium; Calbiochem 344930). LM-211 was purified from the medium with heparin affinity chromatography (heparin agarose, Sigma H6508) and concentrated from the 250 to 500 mM NaCl 50mM Tris-HCl pH7.4 fraction, followed by dialysis in reduced NaCl buffer (90mM).

### Cell culture

293T cells (CRL-3216) were purchased from ATCC and maintained in Dulbecco’s Modified Eagle’s Medium (DMEM; Gibco, # 11885084) supplemented with 10% heat inactivated fetal bovine serum (Gibco, #26140079) and 1% penicillin/streptomycin at 37°C in a humidified 5% CO2 incubator. Neuro-2a cells (CCL-131) were purchased from ATCC and maintained in Minimum Essential Media (Gibco, #11095080) supplemented with 10% heat inactivated fetal bovine serum (Gibco, #26140079) and 1% penicillin/streptomycin at 37°C in a humidified 5% CO2 incubator. C2C12 cells were kindly provided by Clay F. Semenkovich and maintained in DMEM (Gibco, #11965092) supplemented with 10% heat inactivated fetal bovine serum (Gibco, #26140079) and 1% penicillin/streptomycin at 37°C in a humidified 5% CO2 incubator.

### Plasmid transfection and lentiviral infection

Neuro-2a cells were transfected with plasmids (lentiCRISPR v2, lentiGuide-Puro) using Lipofectamine 3000 (Invitrogen, # L3000015), following the manufacturer’s instructions. Briefly, 120,000 – 150,000 cells were seeded in 12-well cell culture plates, and cells were then transfected with 1 µg of total plasmid DNA, 2µL of P3000 and 3µL of Lipofectamine 3000 per well after 20-24 h of culture. For lentivirus production, 5-6×10^5^ 293T cells were seeded in 6-well culture plates and transfected with 1 µg of psPAX2 (Addgene plasmid # 12260; http://n2t.net/addgene:12260; RRID:Addgene_12260), 0.1 µg of pMD2.G (Addgene plasmid # 12259; http://n2t.net/addgene:12259; RRID:Addgene_12259) and 1µg lentiviral vector using Lipofectamine 3000. At 24 h post-transfection, culture media were changed to DMEM supplemented with 30% heat inactivated fetal bovine serum and 1% penicillin/streptomycin. At 48 h post-transfection, culture supernatants were collected and filtered by 0.45 µm PVDF filter unit (Millipore, #SLHVR33RS). For lentiviral transduction, 10^5^ C2C12 cells were seeded into 6-well culture plates. After 24 h, culture media were replaced with lentivirus-containing medium supplemented with 8 µg/mL polybrene (Sigma-Aldrich, # TR-1003). At 24 h post-transduction, C2C12 cells were selected by 300-400 µg/mL hygromycin B (Gibco, #10687010) and 10 µg/mL Puromycin dihydrochloride (Sigma, #P9620).

### T7E1-mediated mismatch cleavage assay

Genomic DNA was extracted from cultured cells using the Quick-DNA Microprep Kit (Zymo Research, #D3020). A 500-600 bp DNA fragment containing the gRNA target site was amplified using GoTaq Flexi DNA polymerase (Promega, #M8295). The amplicons were purified using the QIAquick PCR Purification Kit (QIAGEN, #28106). A total of 500 ng of purified PCR products were denatured at 95°C for 5 minutes and then gradually cooled from 95°C to 25°C over 1.5 hours to form heteroduplexes. The heteroduplexes were then incubated with T7 endonuclease I (NEB, #M0302L) at 37°C for 30 minutes. T7 endonuclease I-mediated mismatch cleavage was analyzed by DNA electrophoresis. The sequence of primers for PCR is listed in Table *S4*.

### Real time quantitative reverse transcription PCR

RNA was extracted from cultured cells using TRIzol reagent (Invitrogen, #15596018) according to the manufacturer’s instructions. Purified RNA was then reverse transcribed into cDNA using PrimeScript RT Master Mix (TaKaRa, #RR036A). For quantitative PCR, Fast SYBR Green Master Mix (Applied Biosystems, #4385612) and a QuantStudio 6 Flex Real-Time PCR System were used, following the manufacturer’s instructions. The sequence of primers for qRT-PCR is listed in Table *S5*.

### RT-PCR

RNA was extracted from kidney cortical tissues using TRIzol reagent (Invitrogen, #15596018) according to the manufacturer’s instructions. Purified RNA was then reverse transcribed into cDNA using PrimeScript RT Master Mix (TaKaRa, #RR036A). For RT-PCR, GoTaq Flexi DNA polymerase (Promega, # M8295) and a C1000 Thermal Cycler were used, following the manufacturer’s instructions. The sequence of primers for RT-PCR is listed in Table *S6*.

### Animal studies

Animal experiments were performed in accordance with the National Institutes of Health Guide for the Care and Use of Laboratory Animals and were approved by the Washington University Institutional Animal Care and Use Committee. *Rosa26*-Cas9 knockin (stock #026179), *Rosa26*-LSL-Cas9 knockin (stock #026175) ^61^, *Igs2*-dCas9-KRAB knockin (stock #030000), *Hopx*-CreERT2 knockin (stock #017606), and Ai14 knockin (stock #007914) ^62^ mice were purchased from the Jackson Laboratory. *Col4a5* (G5X) mutant mice were provided by Yoav Segal, University of Minnesota ^23^. Pax3-Cre Tg ^63^ and Hopx-Cre ^64^ mice were generously provided by Dr. Jonathan Epstein, University of Pennsylvania. Nphs2-Cre transgenic ^65^, *Lamb2* knockout ^24^, and *Cd2ap* knockout ^25^ mice were described previously.

Tg: Lama2-gRNA mice were produced in this study. The transgene was digested with *Kpn*I (NEB, #R3142S) and *Xma*I (NEB, #R0180S), gel-purified to remove plasmid vector sequences using the Wizard SV Gel and PCR Clean-Up System (Promega, #A9281), and further purified by ethanol precipitation. The purified transgene was microinjected into the pronuclei of single-celled B6CBAF2/J embryos by the Washington University Mouse Genetics Core. For recombinant hLM-211 protein injection, mice were anesthetized with isoflurane and injected i.v. with either PBS or hLM-211 (1.2 mg/kg) through the retroorbital sinus for two consecutive days. Kidneys were harvested 24 h after the final injection for histological analysis.

### Immunofluorescence

For immunofluorescence of LAMA2 and COL4A5, fresh tissues were embedded in Tissue-Tek O.C.T. Compound (Sakura Finetek, #4583) without fixation and frozen using liquid nitrogen. For tdTomato and ITGA8, tissues were collected after perfusion fixation with 4% paraformaldehyde/PBS (pH 7.4). The tissues were then immersed in 4% paraformaldehyde/PBS (pH 7.4) for 2 h, followed by cryoprotection in 10% sucrose/PBS and subsequently in 30% sucrose/PBS overnight. The fixed and cryoprotected tissues were embedded in Tissue-Tek O.C.T. Compound and frozen using liquid nitrogen. Nidogen was stained in both unfixed and fixed sections. 7 µm tissue sections were immersed in PBS to remove the O.C.T. Compound and then incubated with 1% BSA/PBS at room temperature for 1 h. The sections were then incubated with primary antibodies in 1% BSA/PBS overnight. After primary antibody incubation, the sections were washed three times with PBS for 5 min each, followed by incubation with secondary antibodies and Hoechst 33258 (Invitrogen, #H3569) at room temperature for 1 h. The sections were then washed with PBS three times for 5 min each and mounted in 90% glycerol in 0.1X PBS containing 1 mg/ml para-phenylenediamine.

Immunofluorescent images were captured using a Nikon C1 confocal system. The primary antibodies used in this study were as follows: polyclonal rabbit anti-Laminin-α2 LG (previously described ^66^, 1:200 dilution), monoclonal rat anti-Nidogen-1 (clone ELM1, Sigma-Aldrich #MAB1946-I, 1:100 dilution), monoclonal rat anti-COL4A5 (Clone H53, Chondrex #7078, 1:200 dilution), polyclonal rabbit anti-pan-Laminin (Sigma Aldrich #L9393, dilution 1: 200), polyclonal goat anti-COL4A1/2 (Southern Biotech #1340-01, 1:200 dilution), and polyclonal goat anti-integrin-α8 (R&D Systems #AF4076, 1:100 dilution). Fluorescent images were acquired on a Nikon C1 confocal microscopy. For near super resolution imaging, images were acquired on a Zeiss LSM 880 Confocal with Airyscan.

### Western blotting

5 µL of plasma from each mouse was diluted in RIPA buffer (0.05 M Tris-HCl [pH 7.5], 0.15 M NaCl, 1% v/v Nonidet P-40, and 1% w/v sodium deoxycholate) and subsequently mixed with 4× Laemmli SDS sample buffer to yield a final 1× concentration. The samples were heated at 100 °C for 10 minutes, then loaded onto a NuPAGE 3–8% Tris-Acetate gel (Invitrogen, #EA0378BOX) for electrophoretic separation. Following gel electrophoresis, the separated proteins were transferred onto a PVDF membrane (Invitrogen, #IB34002) using an iBlot3 Western Blot Transfer System (Invitrogen, #IB31001) with the preset program for high molecular weight proteins. The PVDF membrane was blocked with 5% skim milk in 0.1% PBS-T for 1 hour at room temperature. Thereafter, the membrane was incubated with primary anti-LAMA2 antibody (LG) diluted (1:1000) in CanGet Signal solution 1 (TOYOBO, #NKB-201) overnight at 4 °C, followed by incubation with a secondary HRP-conjugated anti-rabbit antibody diluted (1:2000) in CanGet Signal solution 2 (TOYOBO, #NKB-301) for 1 hour at room temperature. Chemiluminescence imaging was performed using the ECL Select Western Blotting Detection Reagent (Cytiva, #RPN2235) and the iBright FL1500 (Invitrogen, #A44115).

### Statistical analysis

Data in the bar graphs are presented as mean ± standard error, with individual data points shown as dots. One-way ANOVA with Dunnett’s multiple comparisons test was used for multiple group comparisons.

## Acknowledgments

We thank the WashU Mouse Genetics Core for production of Tg: Lama2-gRNA mice, the WashU Center for Cellular Imaging for Super resolution imaging, and Jennifer Richardson for mouse genotyping. We also thank Jonathan Epstein (University of Pennsylvania) for providing transgenic mice. This work was supported by the National Institute of Diabetes and Digestive and Kidney Diseases grant R01DK128660 (J.H.M.), the Children’s Discovery Institute of Washington University and St. Louis Children’s Hospital Pediatric Disease Mouse Models Core (J.H.M.), a grant from the Japan Society for the Promotion of Science Program for Postdoctoral Fellowships for Research Abroad (K.O.), the Cell Science Research Foundation Program for Fellowships for Early Career Researchers B12021-001 (K.O.), JST, ACT-X Grant Number JPMJAX2323, Japan (K.O.), RIKEN intramural funding for Special Postdoctoral Researcher (K.O.), JSPS Kakenhi 23K19359, 24K18132 (K.O.) and MEXT Kakenhi 23H04927, 23H04928 (H.F.).

## Author Contributions

K.O. designed the research, conducted experiments, analyzed data, and wrote the manuscript. M.H.L. conducted experiments, analyzed data, and edited the manuscript. P.P. and K.K.M. conducted experiments and analyzed data. J.H.M. designed the research, analyzed data, and edited the manuscript. P.D.Y. designed the research and analyzed the data. H.F. provided resources and reviewed the manuscript. All authors discussed the results and provided input on the manuscript.

## Competing interests

JHM is a member of the Alport Syndrome Foundation Scientific Advisory Research Network and has served as a consultant for Eloxx Pharmaceuticals, Visterra, and Travere. All other authors declare they have no competing interests.

## Data and materials availability

Data will be made available on request.

---

**Figure S1.**
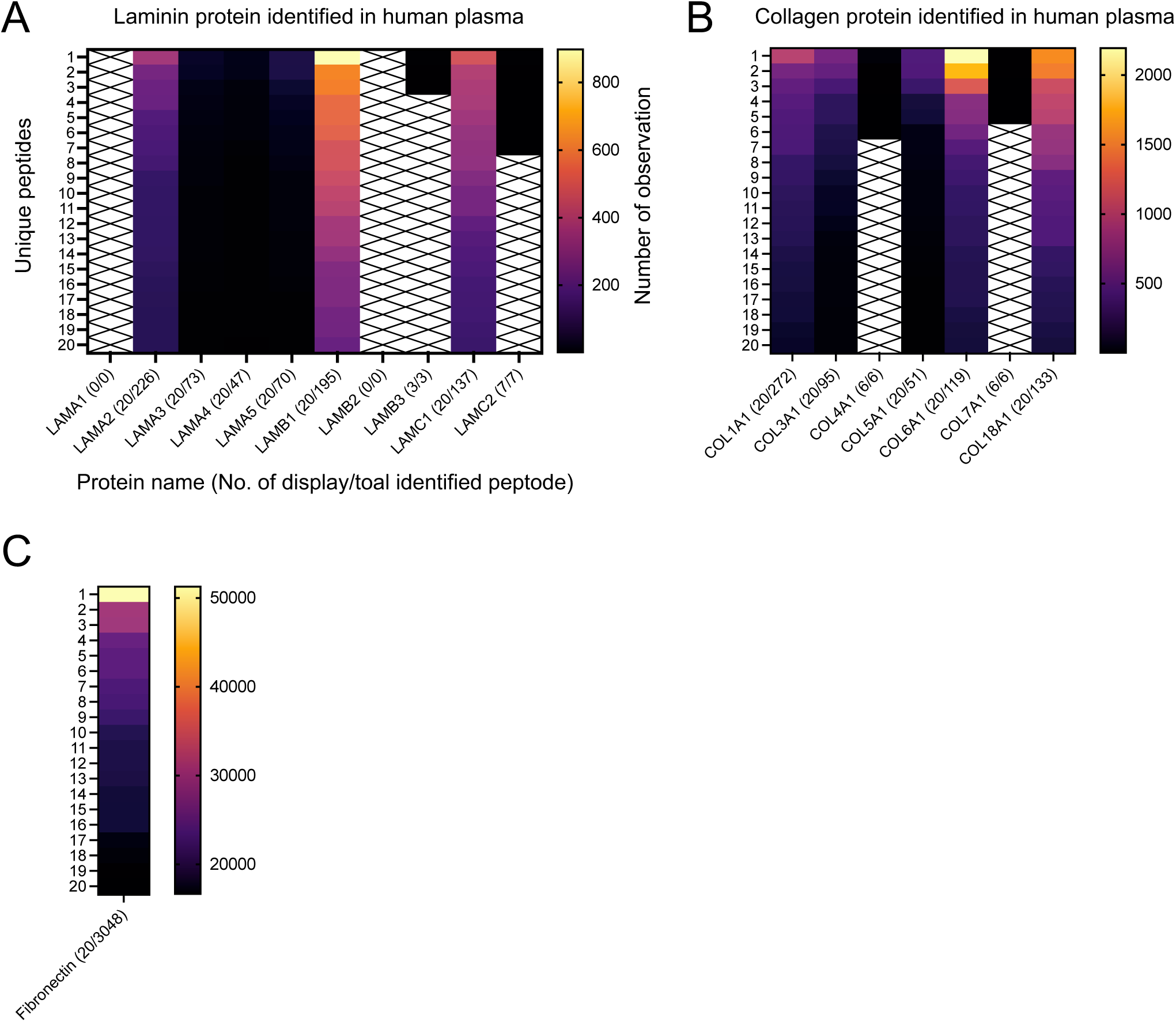
ECM-derived peptides identified in human plasma. (A) Peptides from *LAMA2*, *LAMB1*, and *LAMC1* were more abundant in human plasma compared to other laminin proteins. (B) Collagen peptides, including *COL1A1*, *COL3A1*, *COL5A1*, *COL6A1*, and *COL18A1*, were detected in human plasma. (C) Fibronectin peptides were identified, consistent with previous reports.

**Figure S2.**
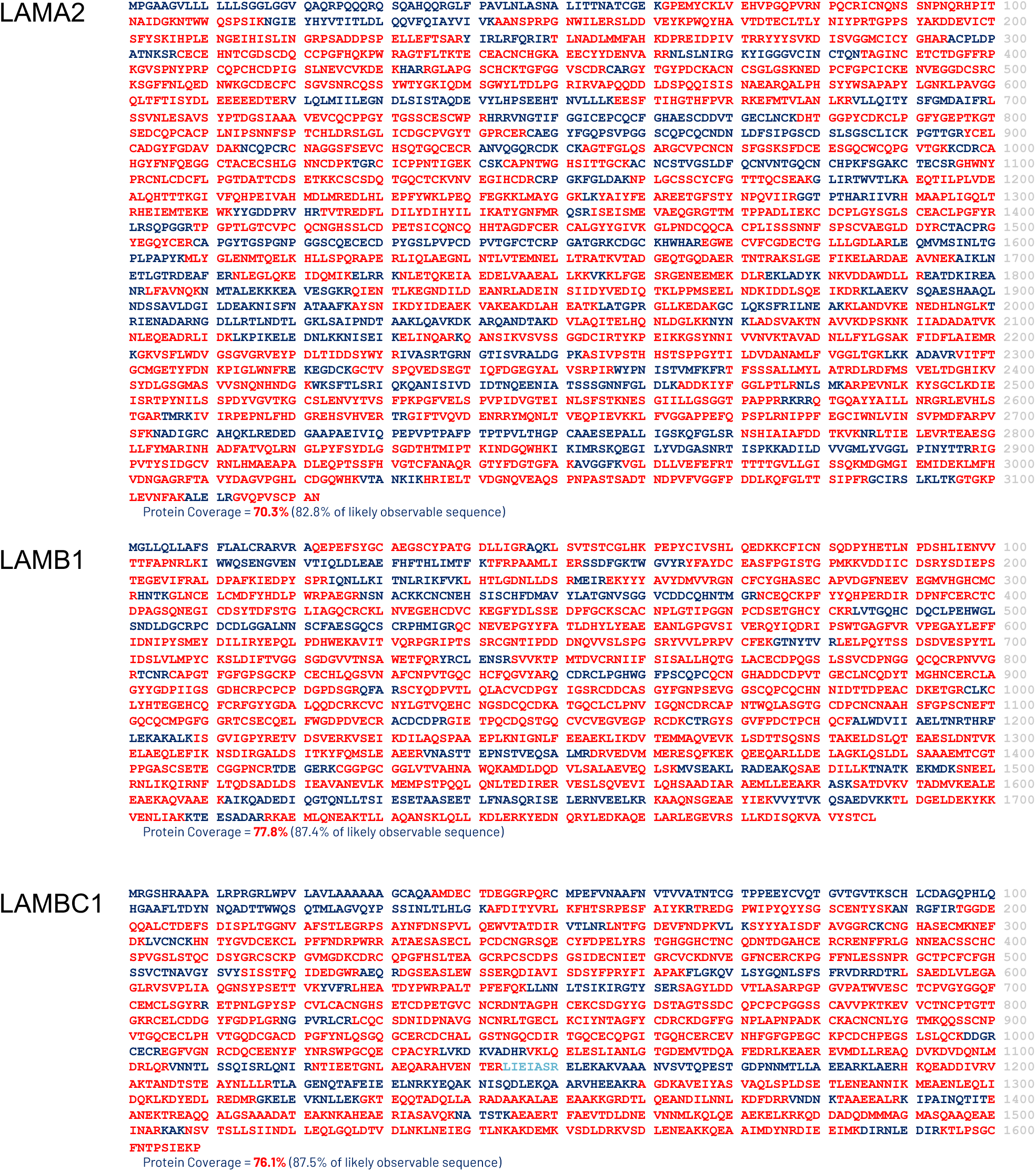
Protein coverage of ECM-derived peptides identified in human plasma for LAMA2, LAMB1, and LAMC1. Peptides identified in human plasma span the entire protein sequences of LAMA2, LAMB1, and LAMC1, with protein coverage rates of 70.3%, 77.8%, and 76.1% (marked in red), respectively.

**Figure S3.**
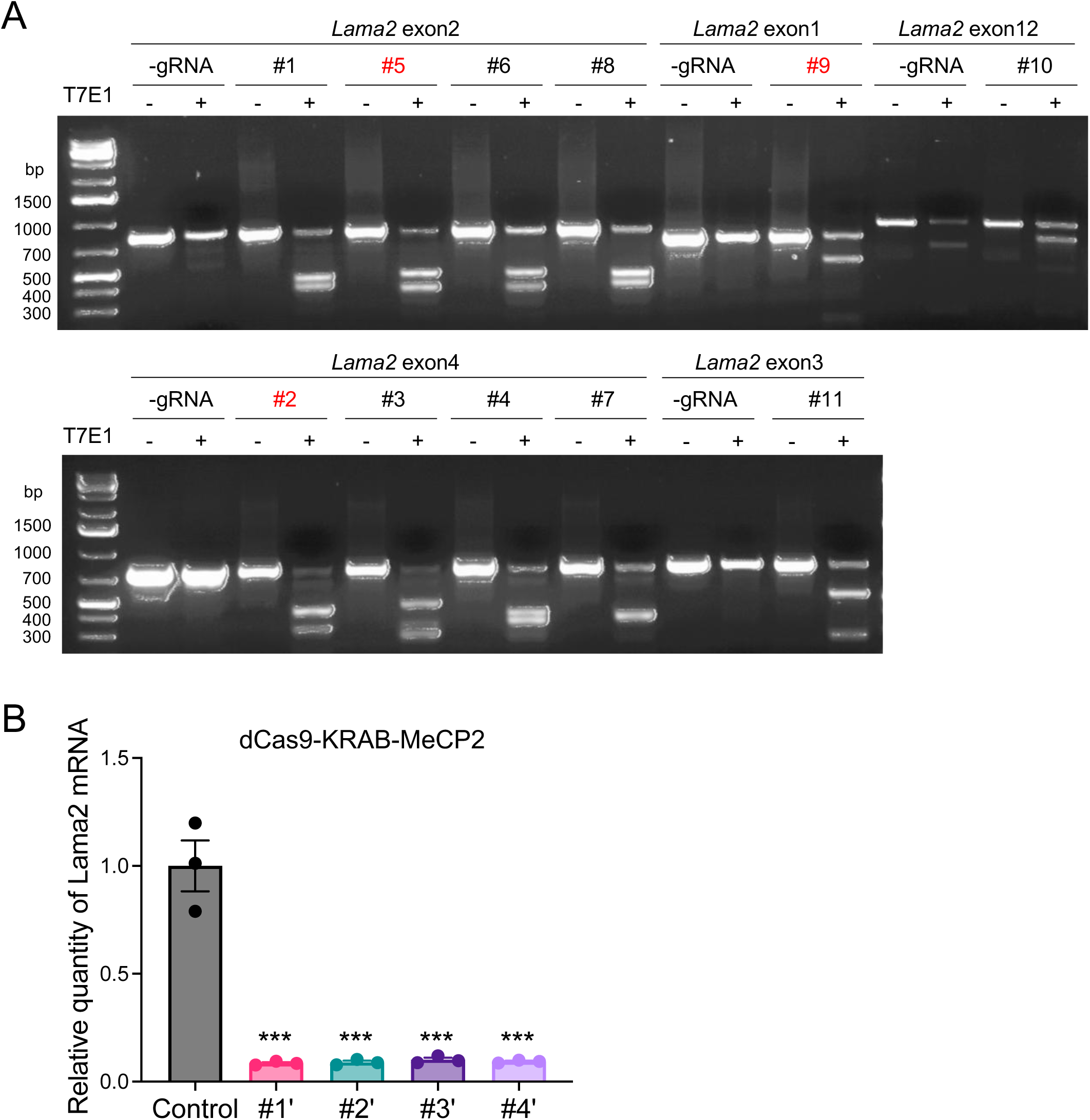
sgRNA screening in cultured cells. (A) T7E1 assay demonstrating the activity of sgRNAs targeting the Lama2 gene in Cas9 and individual sgRNA-transfected Neuro-2a cells. sgRNAs #9, #5, and #2 were selected for use in Tg: Lama2-gRNAs mice. (B) Lama2 transcriptional repression in C2C12 cells mediated by dCas9-KRAB-MeCP2, as assessed by qRT-PCR.

**Figure S4.**
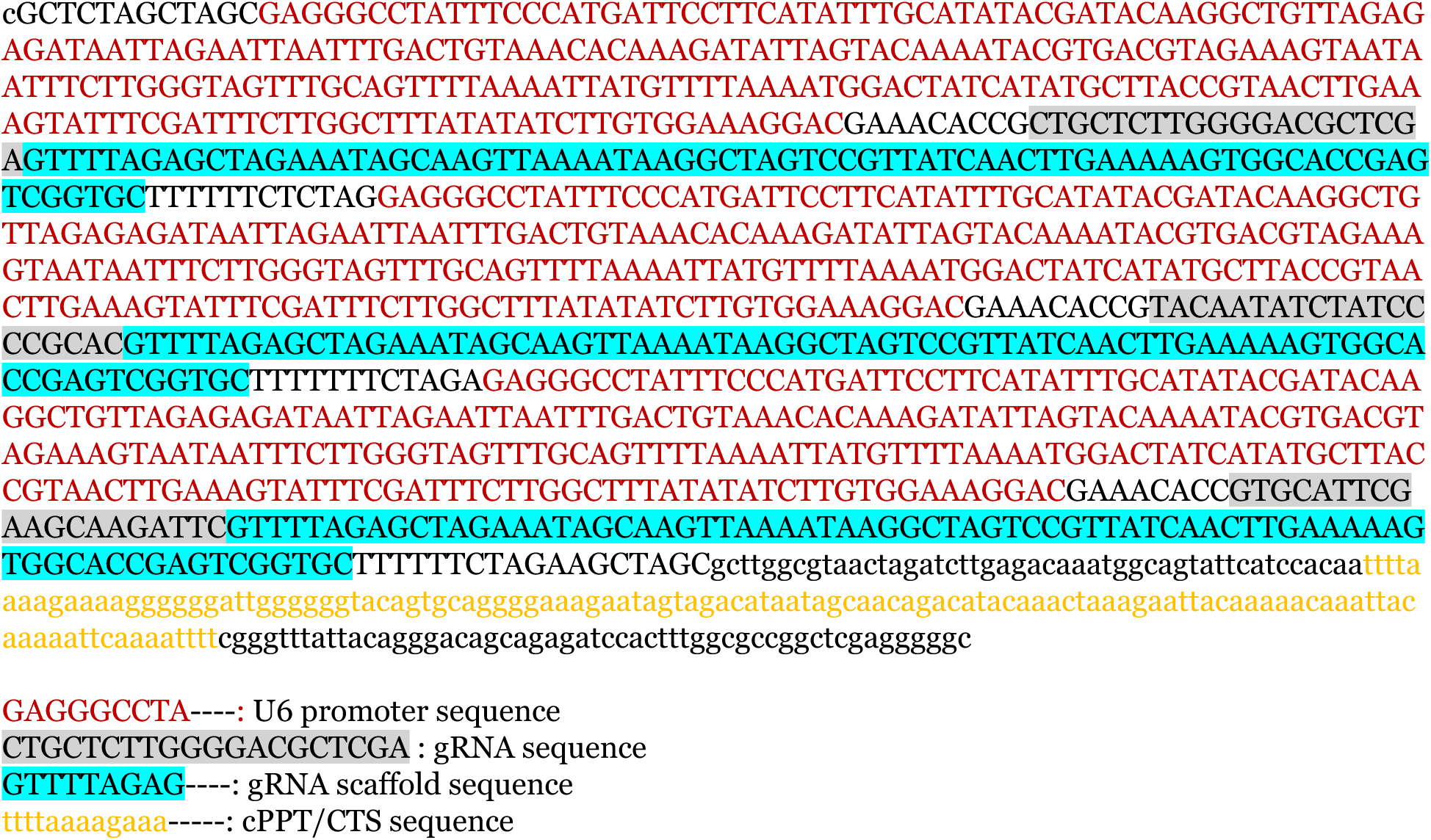
Transgene sequence of Tg: Lama2-gRNAs. The transgene construct consists of three tandemly arrayed [U6 promoter – sgRNA scaffold] cassettes.

**Figure S5.**
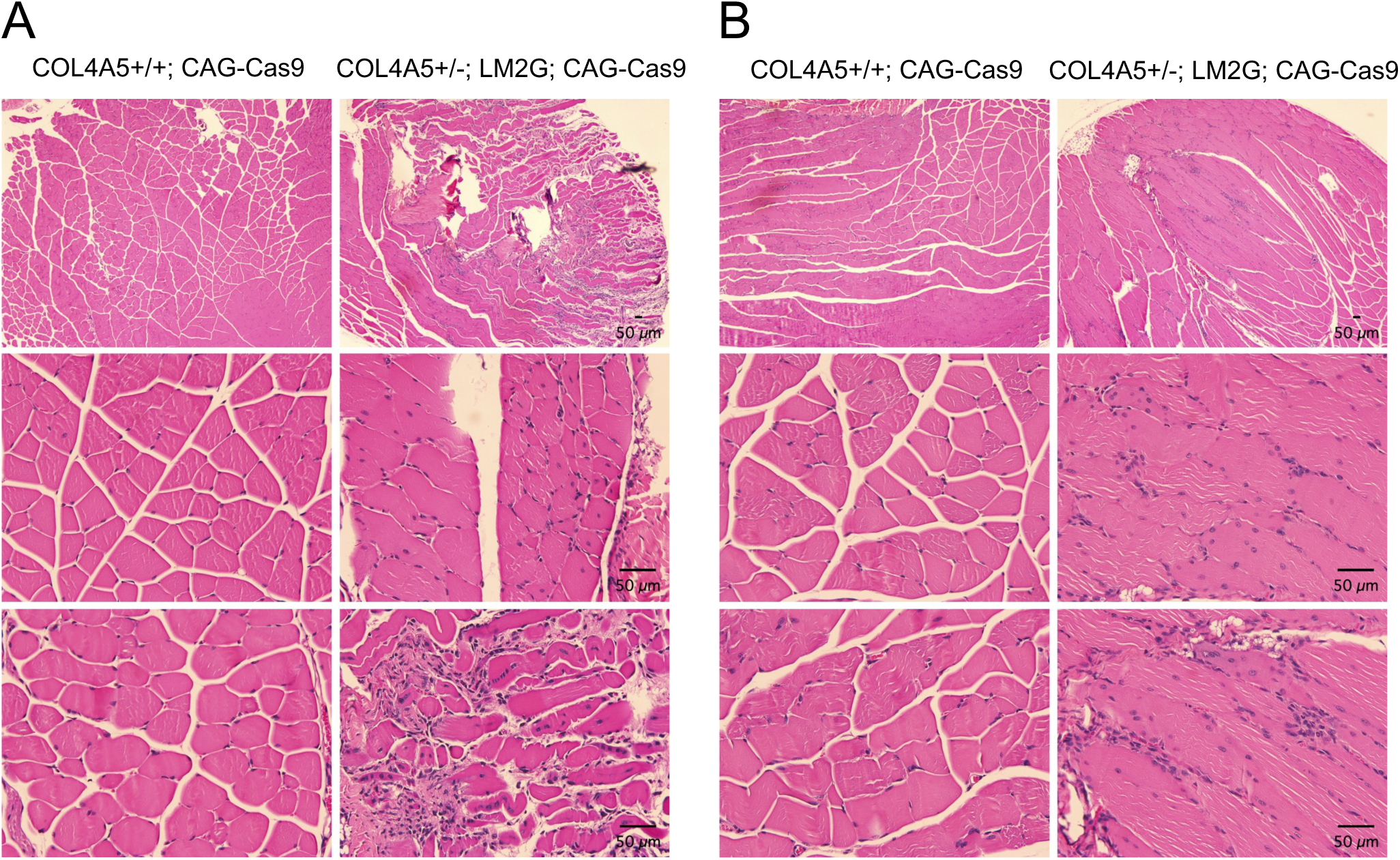
Ubiquitous Cas9-mediated *Lama2* knockout induces muscular dystrophy. Ubiquitous Cas9-mediated disruption of *Lama2* resulted in a muscular dystrophic phenotype, as observed in (A) arm muscle and (B) quadriceps, assessed by H&E staining.

**Figure S6.**
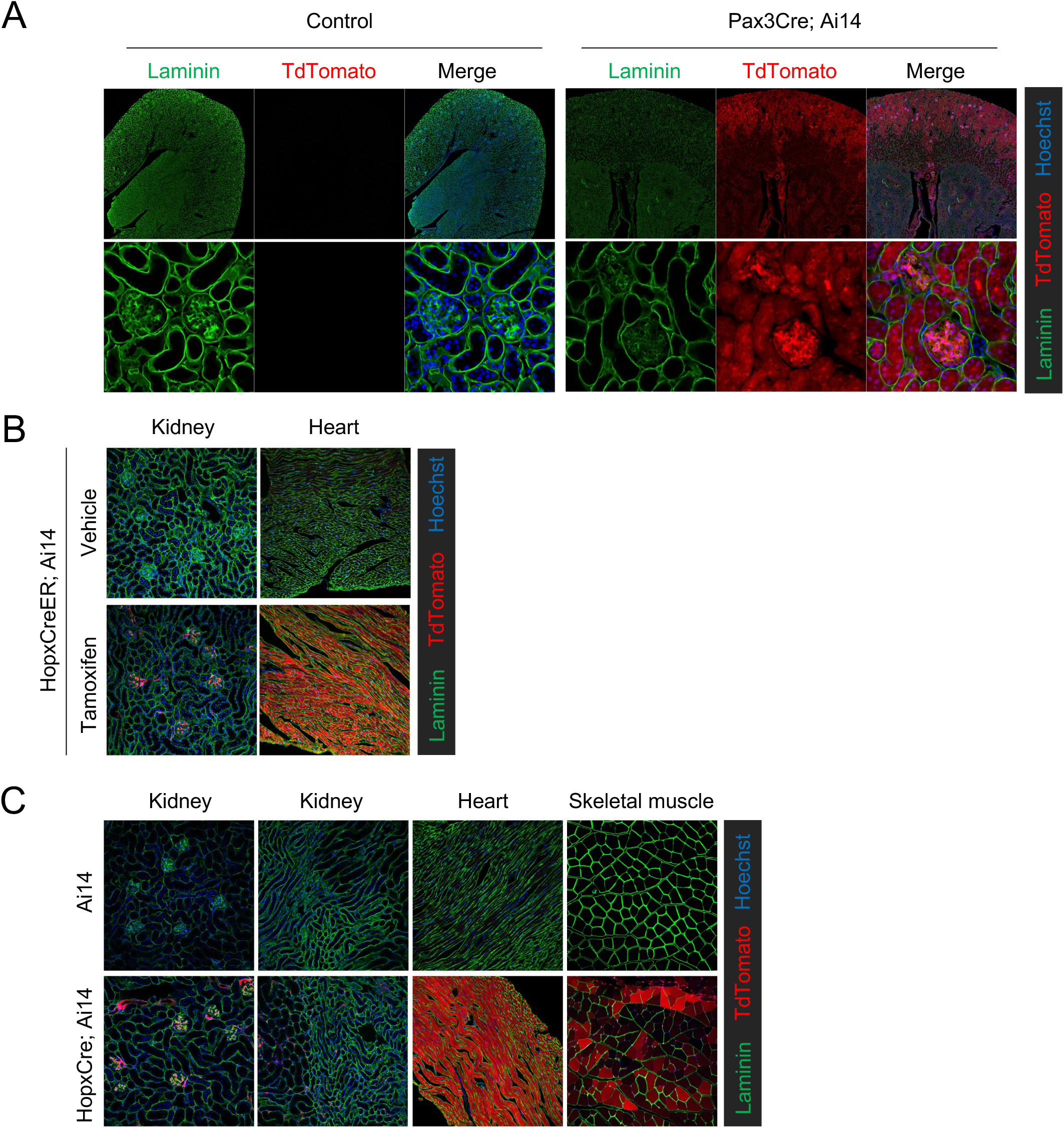
Cell-specific Cre activity assessed by Ai14 reporter expression. (A) *Pax3-Cre*; (B) *Hopx-CreER* with tamoxifen administration; and (C) *Hopx-Cre* induced TdTomato expression in *Ai14*reporter mice. *Pax3-Cre* introduced TdTomato expression in kidney cortical tissue, with nearly all glomerular cells labeled. In *Hopx-CreER; Ai14* and *Hopx-Cre; Ai14* mice, efficient labeling of mesangial cells in the kidney was observed.

**Figure S7.**
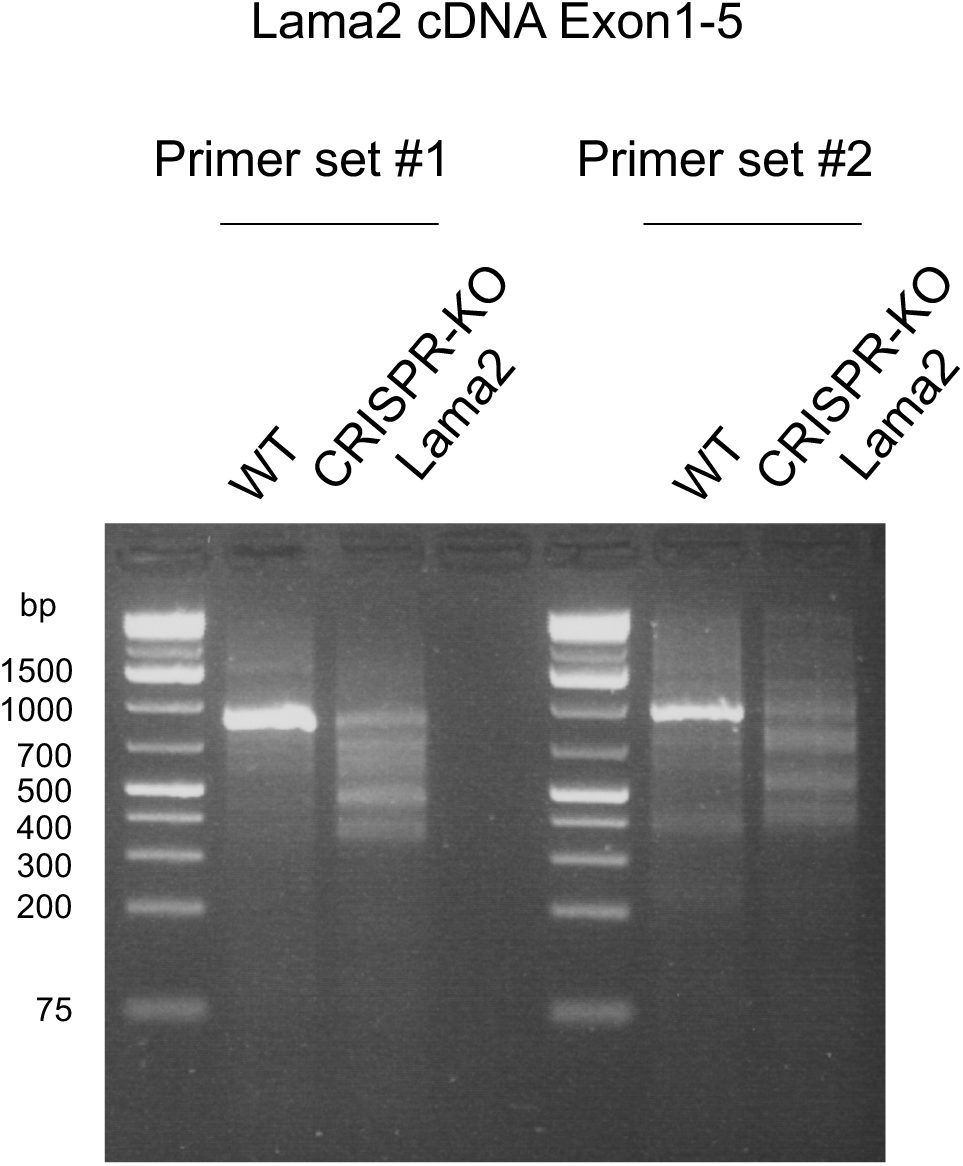
Cas9-mediated disruption of *Lama2* coding sequence. RNA was extracted from kidney cortical tissues of wild-type and *Pax3-Cre; CAG-LSL-Cas9; Tg: Lama2-gRNAs* mice and subjected to RT-PCR. Two distinct primer sets were used to amplify *Lama2* cDNA spanning exons 1–5. *Pax3-Cre; CAG-LSL-Cas9; Tg: Lama2-gRNAs* induced fragmentations in the *Lama2* cDNA sequence across exons 1–5.

**Figure S8.**
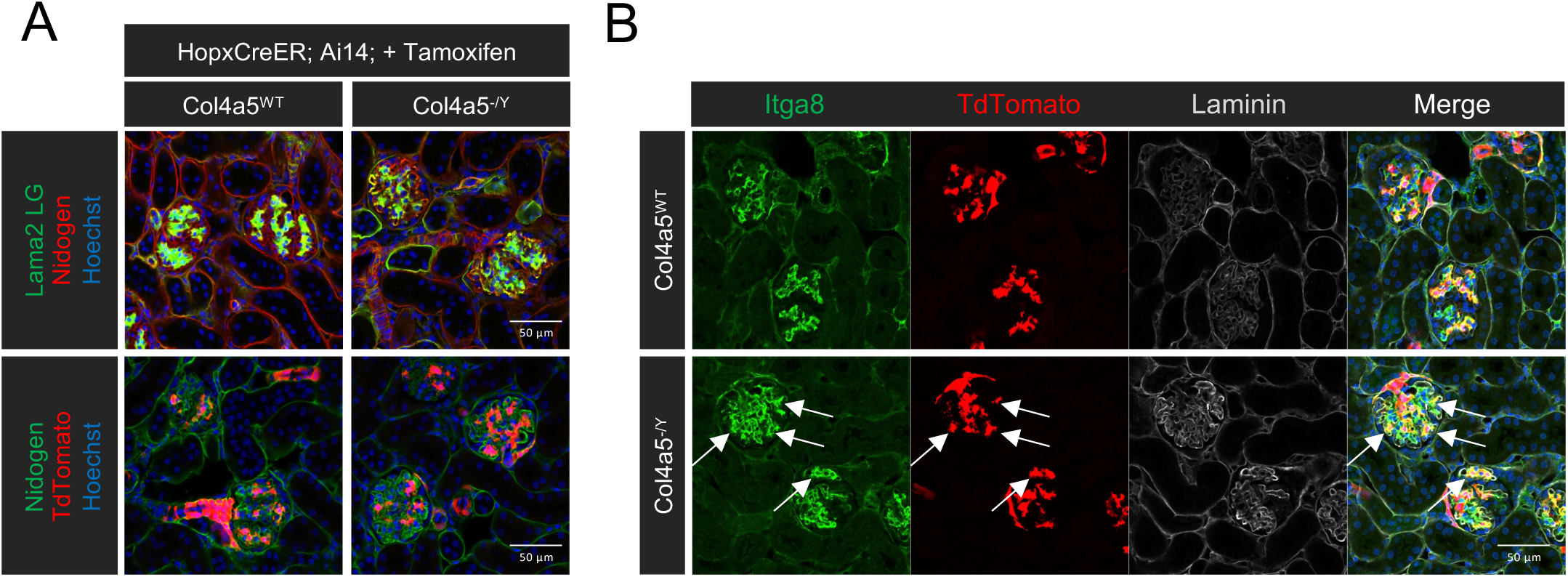
Lack of co-expression of integrin-α8 with TdTomato-positive mesangial Cells. (A) *Hopx-CreER; Ai14* reporter mice reveal no migration of TdTomato-labeled mesangial cells into the LAMA2-positive capillary loops of the glomerular basement membrane (GBM) in Alport syndrome. (B) Immunofluorescence signals of Itga8 and Hopx-TdTomato reporter signals show an expansion of Itga8-positive cells within the Alport glomerulus (arrows), with no colocalization between Itga8 and Hopx-TdTomato in Alport syndrome.

